# Compress global, dilate local: Intentional binding in action–outcome alternations

**DOI:** 10.1101/507582

**Authors:** Shu Imaizumi, Yoshihiko Tanno, Hiroshi Imamizu

**Author notes:** Corresponding author. Present address: Institute for Education and Human Development, Ochanomizu University, Tokyo, Japan.

## Abstract

Intentional binding refers to subjective temporal attraction between an action and its outcome. However, the nature of intentional binding in multiple actions remains unclear. We examined intentional binding in alternated action–outcome dyads. Participants actively or passively pressed a key, followed by a tone, and they again pressed the key; resulting in four keypress–tone dyads in a trial. Participants reproduced the duration of alternated keypress–tone dyads or the temporal interval between a dyad embedded in the alternations. The reproduced duration was shorter in the active than in the passive condition, suggesting the intentional binding in action–outcome alternations. In contrast, the reproduced interval between a dyad was longer in the active condition and did not correlate with the reproduced duration. These results suggest that subjective time during actions relies not only on an internal clock but also on postdictive biases that are switched based on what we recall.

## 1. Introduction

Voluntary motor action is performed for goal-directed behavior and social communications, and concurrently modulates the perception of events in the external world including sensory outcomes of one’s own action. The sensorimotor system in the brain predicts reafferent inputs as sensory outcomes of actions, and consequently modulates the perceptual intensity and neural responses (Blakemore, Wolpert, & Frith, 2000; Weiss, Herwig, & Schutz-Bosbach, 2011). These functions potentially have a role in attributing the source of the sensory outcome to the self, others, or environment, and could be the bases for social interaction. Recent studies have also elucidated that motor action modulates subjective time such as temporal order of events and passage of time flow (Merchant & Yarrow, 2016). For instance, the subjective duration of visual events can dilate during periods of preparation (Hagura, Kanai, Orgs, & Haggard, 2012) and execution of hand movements (Imaizumi & Asai, 2017; Press, Berlot, Bird, Ivry, & Cook, 2014). Dynamic changes in the subjective time in motor action could benefit detection and integration of multiple sensory signals from the environment and motor planning of the subsequent behaviors.

### 1.1. Intentional binding: Compression of action–outcome interval

Different types of time perception bias can occur following the execution of a voluntary, self-paced manual action (e.g., keypress). The timing of the action and its subsequent sensory outcome (e.g., beep tone) can be perceived to shift towards each other (Haggard, Clark, & Kalogeras, 2002). The subjective temporal attraction of an action and its sensory outcome has been called as *intentional binding* (Haggard et al., 2002), although this can also be termed *temporal binding* and *causal binding* because it can also result from causal relation between two successive events even without intentional action (Buehner, 2012; Buehner & Humphreys, 2009). The degree of intentional binding can be an implicit measure of *sense of agency*, referring to a feeling of generating one’s own actions (Gallagher, 2000), and thereby, events in the external world (Haggard & Tsakiris, 2009). Many studies employing the intentional binding paradigm have elucidated the neural, cognitive, emotional, and psychopathological factors underlying sense of agency (Haggard, 2017; Moore & Obhi, 2012).

Intentional binding has been measured by the Libet clock task (Haggard et al., 2002; Libet, Gleason, Wright, & Pearl, 1983; Moore & Haggard, 2008), where participants report the position of the clock hand when they press a key or when the subsequent tone appears. Another established task is the interval estimation task, in which participants verbally report the perceived temporal interval between an action and its outcome (Caspar, Christensen, Cleeremans, & Haggard, 2016; Engbert, Wohlschlager, & Haggard, 2008; Engbert, Wohlschlager, Thomas, & Haggard, 2007; Humphreys & Buehner, 2009). Intentional binding can be evident in conditions with voluntary active movements, but attenuated by involuntary passive movements (Caspar, Cleeremans, & Haggard, 2015; Engbert et al., 2008; Haggard et al., 2002) or with the passive observation of two sensory events (Dewey & Knoblich, 2014; Humphreys & Buehner, 2009; Imaizumi & Tanno, 2019).

From a different viewpoint, intentional binding may also imply that our subjective time is elastic to enable the maintenance of temporal associations between one’s own action and its outcomes by compressing their perceived temporal interval rather than the actual time. Indeed, subjectively and temporally bound action and its outcome can help generate a sense of agency over the action at a retrospective, inferential stage (Imaizumi & Asai, 2015; Synofzik, Vosgerau, & Voss, 2013). Therefore, if a substantial amount of delay is experimentally added to the sensory feedback of bodily movements, the temporal association of the action and its outcome would be violated and sense of agency would be diminished (Franck et al., 2001; Sato & Yasuda, 2005).

### 1.2. Is intentional binding limited to action–outcome dyad?

Although a vast majority of studies on intentional binding have investigated a single action– outcome dyad (Moore & Obhi, 2012), some recent studies suggest that intentional binding can be extended to various ecological situations. This is not surprising given that our everyday physical movements can generate multiple outcomes and they can be executed in response to a sensory event, such as “go” trigger. Indeed, recent studies have demonstrated that intentional binding can emerge for two consecutive outcomes initiated by a single keypress, that is, the perceived timing of both events are attracted towards the action (Ruess, Thomaschke, Haering, Wenke, & Kiesel, 2018), suggesting that intentional binding cannot be limited to a single action–outcome dyad. Furthermore, when an action is triggered by a preceding stimulus in a go/no-go task, the perceived timing of the stimulus is shifted towards the subsequent action (Yabe & Goodale, 2015). This stimulus–action binding and the classical intentional (action–outcome) binding can occur even in stimulus–action–outcome triad (Yabe, Dave, & Goodale, 2017). These findings imply that biased time perception, including the abovementioned two types of temporal binding, could occur in action–outcome alternations (i.e., repetition of action–outcome dyads). This expectation could be natural because, in our daily life, actions and their outcomes are cyclical and/or consecutive, and likely to be embedded in interpersonal settings such as conversation, music, and sports. However, no study to date has examined intentional binding in action–outcome alternations.

### 1.3. The present study

For the first time, to our knowledge, we investigated intentional binding in action–outcome alternations. Specifically, we tested whether subjective compression of the duration of alternated action–outcome dyads occurs analogous to the intentional binding, and whether the intentional binding in an action–outcome dyad within the alternated dyads can accumulate through these alternations and can explain their compressed duration.

In Experiment 1, participants actively or passively made a keypress (i.e., action) followed by a tone (i.e., outcome). The keypress–tone dyad alternated four times in a trial. According to previous studies (e.g., Haggard et al., 2002), three levels of tone delay were employed to ensure that participants tracked the varying subjective time, and to examine the degree of intentional binding modulated by the action–outcome temporal discrepancy. Participants reproduced the perceived duration of the keypress–tone dyads, and the perceived temporal interval between the first keypress–tone dyad or between the last dyad. We hypothesized that, if intentional binding extends to more ecological action–outcome alternations (i.e., repetition of dyads), the duration of the alternations of active movements and outcomes would be perceived as shorter than that in the condition with passive movements. We also expected that, similar to the known intentional binding between an active finger movement and its outcome (Caspar et al., 2015; Engbert et al., 2008; Haggard et al., 2002), the intervals of the first and the last action–outcome dyads embedded within the action–outcome alternations would be perceived as shorter in the condition with active movements than that in the condition with passive movements. Importantly, if the intentional binding in the action–outcome alternations results from the accumulation of the intentional binding in the local action–outcome dyad, there would be a correlation between the degrees of intentional binding in the action–outcome alternations and the local dyads. However, we cannot surely expect that the last dyad interval would be compressed in the condition with active movement, because it might also be affected by the temporal binding between the preceding tone and the subsequent keypress (e.g., Yabe et al., 2017); rather, it might “dilate.”

A follow-up Experiment 2 was conducted to validate the effectiveness of the time reproduction method to detect intentional binding for a subsecond interval and to rule out potentially confounding effects of active movement on emotional states that affect time perception (Droit-Volet, Fayolle, & Gil, 2011; Eberhardt, Kliegl, & Huckauf, 2016) and intentional binding (Christensen, Di Costa, Beck, & Haggard, 2019).

## 2. Experiment 1

### 2.1. Material and methods

#### 2.1.1. Participants

Twenty-four naïve Japanese undergraduates from The University of Tokyo (4 females; mean age 21.3 years, standard deviation (SD) = 1.2) participated in monetary compensation. The sample size was based on previous studies, which showed the effect of active and passive movements on intentional binding (Caspar et al., 2015; Engbert et al., 2008). According to the Flinders Handedness Survey (FLANDERS) (Nicholls, Thomas, Loetscher, & Grimshaw, 2013; Okubo, Suzuki, & Nicholls, 2014), all participants were right handed (median score = 10, range 7–10). They reported that they had normal or corrected-to-normal vision and no history of neurological or psychiatric illness. Written informed consent was obtained from each participant prior to the experiment. This study was approved by the ethics committee of the Graduate School of Arts and Sciences, The University of Tokyo, and was conducted in accordance with the principles of the Declaration of Helsinki.

#### 2.1.2. Apparatus and stimuli

Auditory stimuli were pure tones with a pitch of 440 Hz at a constant comfortable volume and 100-ms duration. Tones were presented to participants and an experimenter through headphones (MDR-Z500, Sony, Tokyo). A fixation cross “+” and time-reproduction cue “○,” both subtending a visual angle of approximately 0.76°, as well as instructions, were presented in white color against a gray background on a 12.5-inch light emitting diode monitor built into a computer (Latitude 12 7000, Dell, Round Rock, Texas), with a refresh rate of 60 Hz. Participants’ responses were collected via an external numeric keypad (BSTK07, Buffalo, Nagoya, Japan). A box-shaped plastic finger holder with a smooth surface and width × depth × height dimensions of 20 × 18 × 20 mm was attached to the Enter key of the keypad. Participants’ right index finger was inserted into this holder and they could perform active pressing of the Enter key. The experimenter could push the top side of the holder without contact with the participants’ finger inside the holder, such that the participants’ finger was passively moved to press and release the Enter key. A black cardboard box with width × depth × height dimensions of 450 × 300 × 110 mm was placed in front of the participants to obstruct the direct view of their and the experimenter’s hands, and the keypad. Stimulus presentation and response collection were controlled by PsychoPy 1.85.1 (Peirce, 2007) running on Windows 7 Professional 64-bit (Microsoft, Redmond, Washington).

#### 2.1.3. Procedures

Experiments were conducted in a quiet, well-lit room. An experimenter wearing headphones sat next to the participant during the experiment. In the active condition, participants voluntarily pressed the Enter key using their right index finger. In the passive condition, the experimenter pushed the participants’ right index finger down and participants were asked to relax their right index finger so that it could be moved upward and downward passively and flexibly.

At the beginning of each trial (Figure 1), the monitor presented a “Press Enter” prompt. Immediately after the pressing of the Enter key, the fixation cross was presented. Moreover, the first tone was presented with three levels of delay (300, 400, or 500 ms) after the first keypress. After the tone, the Enter key was again pressed. The next tone was presented after the delay, whose duration was fixed within each trial and randomly varied within each block. We employed three levels of delay to ensure that the participants tracked the variations in subjective time (Engbert et al., 2007). The levels between 300 and 500 ms were expected to be adequate for the detection of differences between the active-and passive-keypress conditions (Caspar et al., 2015). These keypress–tone alternations were repeated so that there were five keypresses and four tones in each trial. The fixation cross disappeared immediately after the final keypress, and a time-reproduction cue “○” was presented 700–1000 ms (jittered) after the final keypress. Participants were asked to reproduce a designated duration or temporal interval (see the next paragraph). Time reproduction was performed by keeping the zero key pressed using participants’ left index finger. Immediately after the onset of pressing the zero key, the cue was filled in white to inform participants of the initiation of the reproduced duration. No feedback about performance was provided to participants during the experiment. The inter-trial interval was 800 ms.

**Figure 1.**
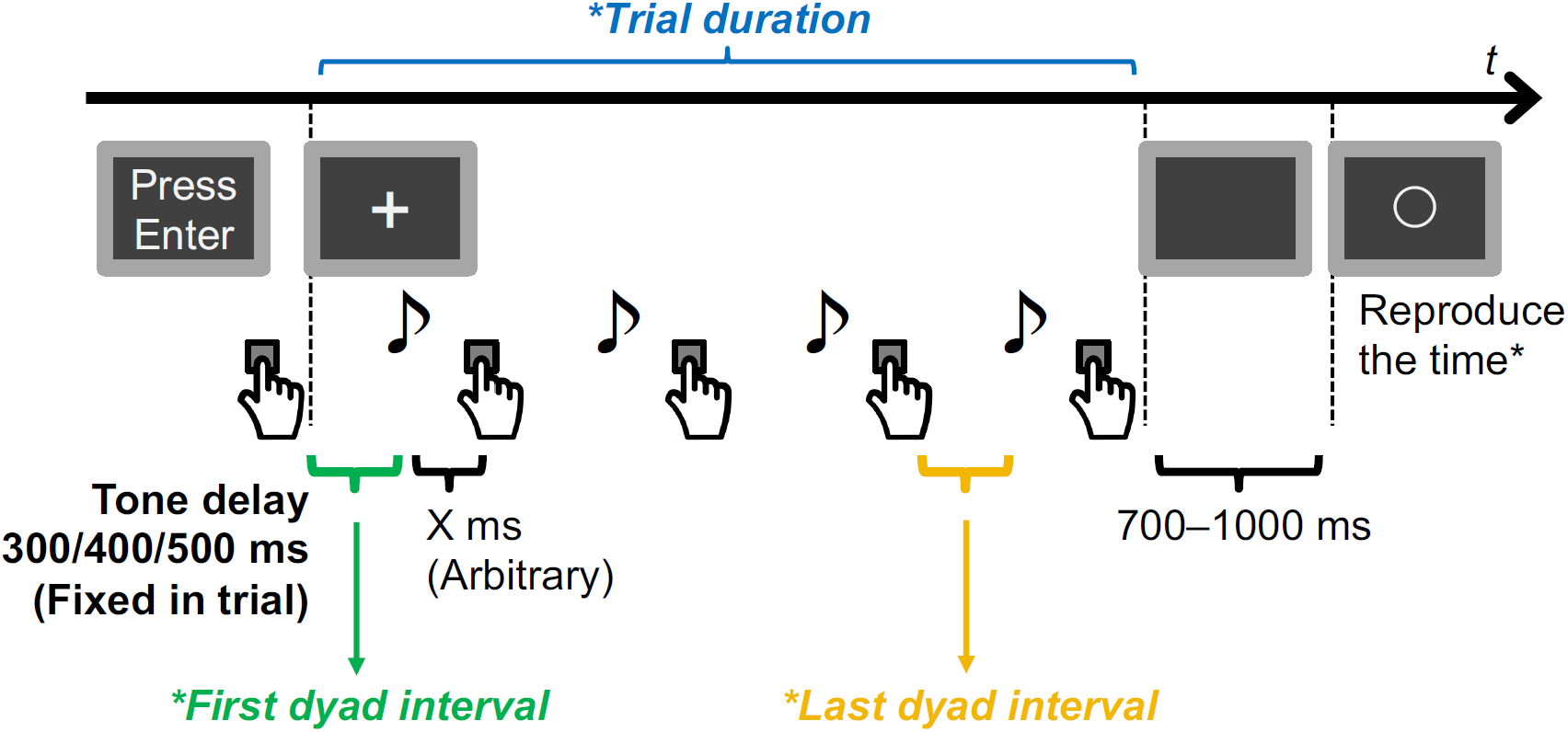
Schematic of the task in Experiment 1. Keypresses were performed by either active or passive finger movement in each block. Participants reproduced one of three types of perceived time (asterisks) in each session.

Participants reproduced the following three types of subjective duration and temporal intervals: the trial duration, first dyad interval, and last dyad interval (Figure 1). The trial duration refers to the subjective duration of the presentation of the fixation cross. The reproduced trial duration was expected to be an index of intentional binding in action–outcome alternations by calculating a differential between the active and passive conditions. The actual trial duration will vary with the lag between a keypress and the preceding tone. The first dyad interval refers to the subjective temporal interval between the first pressing of the Enter key and the subsequent tone, and can be an index of the known intentional binding in an action–outcome dyad (Engbert et al., 2008; Haggard et al., 2002). The last dyad interval refers to the subjective interval between the fourth keypress and the subsequent fifth tone.

Each type of time reproduction was performed in separate session. Each session comprised two blocks each under the active and passive conditions, in an ABAB order. Each block included 15 trials; i.e., three levels of tone delay were repeated five times. The order of the sessions and blocks was counterbalanced across participants. Prior to each session, 12 practice trials (i.e., a trial each for three delays in the four blocks) were performed. Breaks were provided between blocks and sessions, upon request from participants.

#### 2.1.4. Data analysis

For each trial, we calculated the reproduction error by dividing the reproduced duration or interval by the actual value. A larger reproduction error indicates longer perceived duration or interval. We also calculated the coefficient of variation, which is an index of variability of time reproduction, by dividing the SD of the reproduction error by the mean for each participant under each condition.

Reproduction errors and coefficients of variation of the trial duration and the first- and last-dyad intervals were examined using a repeated measures analysis of variance (ANOVA) with two within-factors of *delay* (300, 400, and 500 ms) and *movement* (active and passive). When the sphericity assumption was violated, the Greenhouse–Geisser correction was applied to the degree of freedom for the ANOVA. Post-hoc analyses were performed with Bonferroni correction. Smaller reproduction errors in the active condition than in the passive condition was considered indicative of intentional binding (e.g., Engbert et al., 2008). Finally, we analyzed inter-individual correlations between intentional bindings in the trial duration, and the first- and last-dyad intervals, using the differentials in reproduction error (collapsed across three delay conditions) between the active and passive conditions, using two-tailed test. Analyses were performed using SPSS 24.0 (IBM, Armonk, New York).

### 2.2. Results and discussion

Trials in which any temporal interval between keypress and its preceding tone or reproduced time was out of the range of the individual mean ± 3 SD were excluded from the analyses. As a result, for the session measuring the trial duration, a median 1.67% of the trials were excluded (interquartile range (IQR): 1.67–3.33 %). For the sessions measuring the first- and last-dyad intervals, a median 1.67% (IQR 0.00–1.67%) and 1.67% (IQR 1.25–2.08%) of the trials were excluded, respectively.

#### 2.2.1. Trial duration

The actual trial durations in the 300-, 400-, and 500-ms delay conditions were 2185 ± 179 ms, 2526 ± 161 ms, and 2901 ± 167 ms (mean ± SD), respectively. As shown in Figure 2A, we found a significant main effect of delay on the reproduction error of the duration of action–outcome alternations (*F*(1.52, 34.98) = 28.92, *p* < .001, *η*^2^_p_ = .557) indicating a greater overestimation in the condition with a shorter delay. This could be interpreted as a central tendency (Hollingworth, 1910) where individuals’ estimation of quantities, such as time (Lejeune & Wearden, 2009), tends to gravitate to a representative value (e.g., mean), that is, shorter (longer) durations are perceived as longer (shorter) than they actually are.

**Figure 2.**
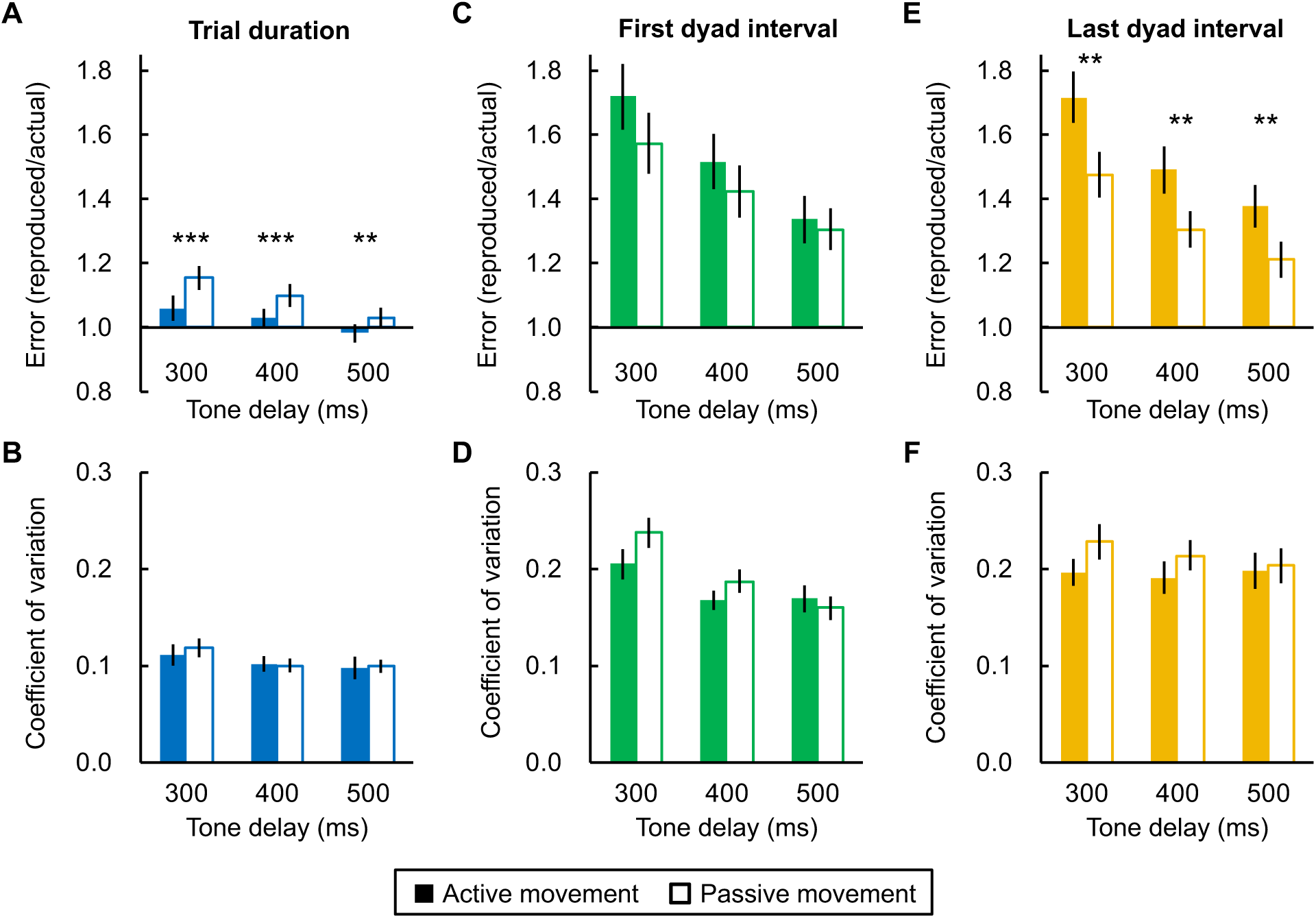
Error and its coefficient of variation for the reproduction of (A, B) the trial duration, (C, D) the first dyad interval, and (E, F) the last dyad interval in Experiment 1. Error bars denote one standard error of the mean. Asterisks represent significant differences between active- and passive-movement conditions (^***^*p* < .001, ^**^*p* < .01: Bonferroni-corrected).

There was also a significant main effect of movement on the reproduction error (*F*(1, 23) = 35.11, *p* < .001, *η*^2^_p_ = .604), indicating a shorter reproduced duration in the active condition than in the passive condition, that is, an intentional binding-like subjective duration compression. We also found a significant interaction between the two factors (*F*(1.82, 41.94) = 3.99, *p* = .029, *η*^2^_p_ = .148). Simple main effects of delay were found for both movement conditions (active: *F*(2, 22) = 13.37, *p* < .001, *η*^2^_p_ = .549; passive: *F*(2, 22) = 17.57, *p* < .001, *η*^2^_p_ = .615), indicating that a comparable level of the central tendency was found regardless of active and passive movements. Simple main effects of movement were also found for all delay conditions (300 ms: *F*(1, 23) = 44.78, *p* < .001, *η*^2^_p_ = .661; 400 ms: *F*(1, 23) = 24.12, *p* < .001, *η*^2^_p_ = .512; 500 ms: *F*(1, 23) = 9.62, *p* = .005, *η*^2^_p_ = .295). However, smaller effect sizes in the larger delay conditions suggested that those intentional binding-like effects were reduced by the action–outcome temporal discrepancy violating internal prediction (Haggard et al., 2002; Imaizumi & Tanno, 2019; Ruess, Thomaschke, & Kiesel, 2018). This finding was supported by a follow-up ANOVA on the differential between the active and passive conditions, which showed a significant effect of delay (*F*(2, 46) = 3.99, *p* = .025, *η*^2^_p_ = .148) with a significant difference between the 300- and 500-ms delay conditions suggesting a weaker intentional binding for a large delay (*p* = .021; no differences between the other pairs: *p*s > .441).

As for the coefficient of variation (Figure 2B), we found a weak but significant main effect of delay (*F*(2, 46) = 3.34, *p* = .044, *η*^2^_p_ = .127), but post-hoc comparisons suggested no differences between delay conditions (300 ms and 400 ms: *p* = .222; 300 ms and 500 ms: *p* = .134; 400 ms and 500 ms: *p* = .999). Importantly, there was no main effect of movement (*F*(1, 23) = 1.41, *p* = .711, *η*^2^_p_ = .006) nor an interaction (*F*(2, 46) = 0.21, *p* = .810, *η*^2^_p_ = .009), indicating that the variability in time reproduction was not affected by active and passive movements regardless of the amount of tone delay. These suggest that the reproduction error biased by active and passive movements were not merely explained by the variability in time reproduction. In sum, these results suggested that the intentional binding induced by active movements can be observed not only in a single action–outcome dyad, which has been traditionally studied, but also in action– outcome alternations.

#### 2.2.2. First dyad interval

As shown in Figure 2C, there was a significant main effect of delay on the reproduction error of the first action–outcome dyad interval (*F*(1.34, 30.79) = 40.60, *p* < .001, *η*^2^_p_ = .638). Post-hoc comparisons revealed significant differences between all adjacent delay conditions (*p*s < .001), again indicating the central tendency similar to that observed in the reproduction error of the trial duration. There was no main effect of movement (*F*(1, 23) = 2.21, *p* = .151, *η*^2^_p_ = .088) nor that of the interaction between the two factors (*F*(1.29, 29.70) = 2.22, *p* = .142, *η*^2^_p_ = .088), suggesting a lack of intentional binding. A *post hoc* power analysis using G*Power 3.1.9.3 (Faul, Erdfelder, Lang, & Buchner, 2007) indicated that the actual power to find a main effect of movement was .937. Thus, the null effect of movement cannot merely be attributed to the statistical underpower.

As for the coefficient of variation (Figure 2D), we found a main effect of delay (*F*(2, 46) = 10.31, *p* < .001, *η*^2^_p_ = .309) and an interaction with movement (*F*(2, 46) = 3.83, *p* = .029, *η*^2^_p_ = .143). A significant simple main effect of delay was found in the passive condition (*F*(2, 22) = 17.15, *p* < .001, *η*^2^_p_ = .609) but not in the active condition (*F*(2, 22) = 2.61, *p* = .096, *η*^2^_p_ = .192), indicating that the coefficient of variation decreased with increasing tone delay only in the passive condition. Importantly, there was no main effect of movement (*F*(1, 22) = 2.67, *p* = .116, *η*^2^_p_ = .104), suggesting that variability of interval reproduction *per se* cannot explain the lack of intentional binding.

#### 2.2.3. Last dyad interval

As shown in Figure 2E, we found a significant main effect of delay on the reproduction error of the last action–outcome dyad interval (*F*(1.60, 36.67) = 51.40, *p* < .001, *η*^2^_p_ = .691). Post-hoc comparisons revealed significant differences between all adjacent delay conditions (*p*s < .001), again reflecting the central tendency. Notably, we found a main effect of movement (*F*(1, 22) = 9.76, *p* = .005, *η*^2^_p_ = .298), without an interaction effect (*F*(1.37, 31.41) = 1.17, *p* = .306, *η*^2^_p_ = .048), indicating a *longer* perceived interval in the active condition than in the passive condition. As for the coefficient of variation (Figure 2F), there were neither main effects of delay (*F*(2, 46) = 0.81, *p* = .452, *η*^2^_p_ = .034) and movement (*F*(1, 23) = 2.32, *p* = .142, *η*^2^_p_ = .091) nor an interaction between them (*F*(2, 46) = 0.95, *p* = .395, *η*^2^_p_ = .040), indicating that the variability of time reproduction was unaffected by experimental manipulations and did not merely explain the modulated reproduction error. These results suggest that a biased time perception in the direction opposite to the intentional binding was induced by active movement in the final dyad in action– outcome alternations.

#### 2.2.4. Correlations among three measures

We examined correlations between the active movement-induced time distortions in the duration of overall action–outcome alternations, the first action–outcome dyad interval, and the last dyad interval. However, there were no significant correlations between the three measures (trial duration and first dyad interval: *r*(22) = −.136, *p* = .527; trial duration and last dyad interval: *r*(22) = −.091, *p* = .673; first- and last-dyad intervals: *r*(22) = .317, *p* = .131). These results suggest that the compression of perceived duration of the action–outcome alternations induced by active movement cannot merely be explained by the local action–outcome dyad, either at the early or last stage of the trial. The compression of trial duration, the dilation of last dyad interval, and the unaffected first dyad interval may be explained by different mechanisms.

#### 2.2.5. Summary of Experiment 1

Results of Experiment 1 suggested that, analogous to the intentional binding (e.g., Engbert et al., 2008), alternations of active movements and their auditory outcomes are also capable of inducing compression of the subjective duration of the alternations, as compared to those in the condition involving passive movements. Contrary to our expectation, we did not find any biased interval perception for the first action–outcome dyad in the alternations, and found an “dilation” of the interval for the last action–outcome dyad. These biases in time perception did not correlate with each other, which suggested different underlying mechanisms.

Given the complicated nature of our task including multiple keypresses and tones, one may doubt whether participants indeed reproduced temporal interval of the designated action–outcome dyad. For example, if participants reproduced the interval between the first tone and the subsequent second keypress (i.e., mean response time of 234.2 ms across conditions and participants, SD = 35.4) instead of the first dyad interval, and the interval between the preceding third tone and the fourth keypress (i.e., mean response time of 228.7 ms, SD = 37.0) instead of the last dyad interval, the reproduction error would positively correlate with the response time. However, there was no significant correlation among reproduction errors and response times for both the first and last dyad intervals in each condition (*r*s(22) = −.154 to .355; *p*s = .089 to .492, two-tailed test). This suggests that we indeed measured the perceived interval of the first and last dyads.

Before a detailed discussion on the present results, we will address two potential confounding factors in this experiment. First, our findings from contrast between active and passive finger movements might be explained merely by the emotional states (e.g., mood change) induced by the active and passive movements, not by the motor command and voluntariness in active movement themselves. Indeed, experimentally manipulated emotional states can affect supra- and sub-seconds duration judgment (Droit-Volet et al., 2011; Eberhardt et al., 2016). Moreover, the emotional valence of the outcome of an action can affect the degree of intentional binding (Takahata et al., 2012; Yoshie & Haggard, 2013, 2017), and crucially, negative (i.e., fearful, angry) states can also reduce intentional binding (Christensen et al., 2019).

Second, the null effect of active and passive movements on the first dyad interval might be due to the characteristics of the task that measured perceived time. While the time reproduction method used in the present study may provide more accurate measurement than a verbal estimation (Mioni, 2018), which has been used in previous studies on intentional binding (e.g., Engbert et al., 2007), the reproduction of subsecond intervals (around 400 ms in our experiment) may be difficult and too variable given the motor latency required for the reproduction keypresses (van Volkinburg & Balsam, 2014). Furthermore, while intentional binding for a single action– outcome dyad with temporal interval longer than 500 ms has been reliably detected by the reproduction method (Dewey & Knoblich, 2014; Howard, Edwards, & Bayliss, 2016; Humphreys & Buehner, 2010; Poonian & Cunnington, 2013), there is little evidence on whether the reproduction method can accurately detect the intentional binding with the interval shorter than 500 ms (but see Dewey & Knoblich, 2014). Therefore, we conducted a follow-up Experiment 2 to test whether intentional binding for the subsecond interval of an action–outcome dyad could be detected by the time reproduction method, and whether the biased time perception could be explained merely by the mood changes induced by active and passive movements. In a trial, participants performed active or passive keypress, and after the three levels of delay (300–500 ms), they listened to a tone. The temporal interval between the keypress and the tone was reproduced. Participants’ moods were assessed by a questionnaire before and after the task.

## 3. Experiment 2

### 3.1. Material and methods

#### 3.1.1. Participants

New 23 naïve Japanese undergraduates from The University of Tokyo participated for monetary compensation. One participant was excluded from the analysis because her coefficient of variation in the interval reproduction (collapsed across conditions) exceeded mean + 3 SD. Thus, data from 22 participants were analyzed (11 females; mean age 19.0 years, SD = 0.8). They were right-handed (FLANDERS median score = 10, ranged 7–10), and they reported that they had normal or corrected-to-normal vision and no history of neurological or psychiatric illness.

#### 3.1.2. Apparatus and stimuli

The experimental material were identical to those used in Experiment 1, except for a 14-inch light emitting diode monitor built into a computer (Inspiron 14 7000, Dell, Round Rock, Texas) and a custom program written in Hot Soup Processor 3.4 (ONION Software, Japan) running on Windows 10 Professional 64-bit (Microsoft, Redmond, Washington) for stimulus presentation and response collection.

#### 3.1.3. Questionnaire

Participants’ emotional states (i.e., moods) were assessed by the Japanese version of the Positive and Negative Affect Schedule (PANAS) consisting of the Positive Affect and Negative Affect subscales (Sato & Yasuda, 2001; Watson, Clark, & Tellegen, 1988). Each subscale includes eight items rated on a 6-point scale, with a score ranging from 8 to 48. Larger values indicate a stronger positive or negative mood, respectively. The PANAS was completed through the Google Forms tool (Google, Mountain View, California) using iPad Air 2 (Apple, Cupertino, California). The item order was randomized in each administration.

#### 3.1.4. Procedures

Experimental tasks were identical to those in Experiment 1, except for the following (Figure 3A). At the beginning of each trial, the monitor presented the fixation cross followed by the pressing of the Enter key. The subsequent tone was presented 300, 400, or 500 ms after the keypress. The time-reproduction cue was presented 700–1000 ms after the tone. Participants were asked to reproduce the subjective temporal interval between the keypress and the tone. The inter-trial interval was 800 ms.

**Figure 3.**
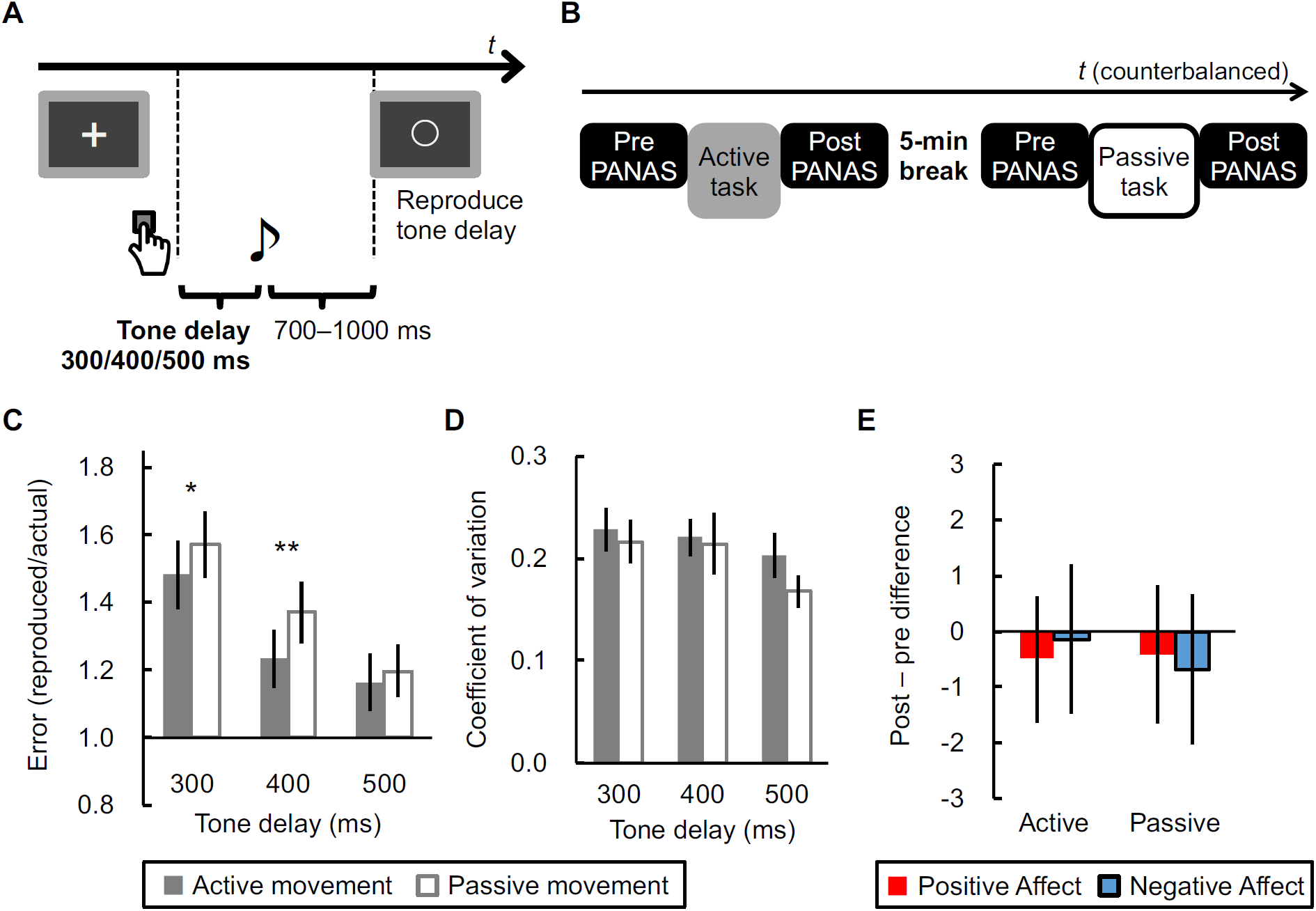
(A) Schematic of the task in Experiment 2. Keypresses were performed by either active or passive finger movement. (B) General procedure of Experiment 2. (C) Error and (D) its coefficient of variation for the reproduction of the tone delay. Error bars denote standard error of the mean. Asterisks represent significant differences between active- and passive-movement conditions (^**^*p* < .01, ^*^*p* < .05: Bonferroni-corrected). (E) Mean differential scores on the subscales (Positive Affect, Negative Affect) of the Positive and Negative Affect Schedule (PANAS) between post- and pre-task administrations. Error bars denote 95% confidence interval.

An overview of the experiment is presented in Figure 3B. Each of the two blocks included ten trials each for the three levels of tone delay under either the active or passive condition. The block order was counterbalanced across participants and the trial order was randomized. Before and after each block, participants completed the PANAS. A five-minute break was provided after the PANAS administration following the first block to eliminate the potential carry-over effect on moods. Participants were instructed to sit relaxed but not to sleep during the break. Prior to the experiment, nine practice trials (three delay conditions were repeated three times) were performed under each of the active and passive conditions.

#### 3.1.5. Data analysis

We calculated the reproduction error by dividing the reproduced interval by the actual value for each trial, and the coefficient of variation, for each participant under each condition. Reproduction errors and coefficients of variation were analyzed using a repeated measures ANOVA with two within-factors of *delay* (300, 400, and 500 ms) and *movement* (active and passive). Sphericity violation and multiple comparisons were treated as in Experiment 1, where appropriate. As for PANAS, differentials of post-task minus pre-task administration for Positive Affect and Negative Affect subscale scores were calculated for each of the active and passive movement conditions.

To check the mood changes induced by the task, the differential scores were tested by a two-tailed one-sample *t*-test against zero. Differential scores were also submitted to an ANOVA with two within-factors of *valence* (Positive Affect and Negative Affect) and *movement* (active and passive). Finally, we analyzed correlations between the degree of intentional binding (i.e., subtraction of the reproduction error in the active condition from that in the passive condition) and the post-pre differentials in the Positive Affect and Negative Affect scores in the active and passive conditions using two-tailed test, to further examine any influence of moods on intentional binding.

### 3.2. Results and discussion

#### 3.2.1. Time reproduction error

Trials with a reproduced interval outside the individual mean ± 3 SD were excluded from the analyses. A median of 1.70% of the trials were excluded (IQR: 1.70–2.90 %). Results have been summarized in Figure 3C. We found significant main effects of delay (*F*(1.53, 32.20) = 57.42, *p* < .001, *η*^2^_p_ = .732) and movement (*F*(1, 21) = 9.42, *p* = .006, *η*^2^_p_ = .310), and their interaction (*F*(2, 42) = 4.48, *p* = .017, *η*^2^_p_ = .176). Simple main effects of delay were found for both movement conditions (Active: *F*(2, 20) = 36.90, *p* < .001, *η*^2^_p_ = .787; Passive: *F*(2, 20) = 40.19, *p* < .001, *η*^2^_p_ = .801), again suggesting a central tendency of time perception as that observed in Experiment 1. Simple main effects of movement were found for the 300-ms (*F*(1, 21) = 5.73, *p* = .026, *η*^2^_p_ = .214) and 400-ms delay conditions (*F*(1, 21) = 14.45, *p* = .001, *η*^2^_p_ = .408) but not for the 500-ms condition (*F*(1, 21) = 1.27, *p* = .273, *η*^2^_p_ = .057), indicating that a shorter perceived interval resulted from active movement only under conditions with a small tone delay. These results reflect the typical characteristics of intentional binding (Haggard et al., 2002; Imaizumi & Tanno, 2019; Ruess, Thomaschke, & Kiesel, 2018).

We did not find any significant main effects of delay (*F*(2, 42) = 2.50, *p* = .094, *η*^2^_p_ = .106) and movement (*F*(1, 21) = 0.82, *p* = .375, *η*^2^_p_ = .038), and their interaction (*F*(2, 42) = 0.56, *p* = .578, *η*^2^_p_ = .026) on the coefficients of variation (Figure 3D). This suggested that the replicated intentional binding cannot be explained merely by the variability in the time reproduction affected by active and passive movements and action-outcome delays. Thus, we concluded that the time reproduction method was able to detect intentional binding for the subsecond scale of the action– outcome interval.

#### 3.2.2. Positive and Negative Affect Schedule (PANAS)

Results of the PANAS have been summarized in Table 1 and Figure 3E. One sample *t*-tests against zero on the post-pre PANAS differentials indicated no significant differences (Positive Affect, active: *t*(21) = −0.87, *p* = .396, *d* = −0.19; Negative Affect, active: *t*(21) = −0.20, *p* = .843, *d* = −0.04; Positive Affect, passive: *t*(21) = −0.65, *p* = .523, *d* = −0.14; Negative Affect, passive: *t*(21) = −1.00, *p* = .331, *d* = −0.21), suggesting that our experimental task with active and passive finger movements did not affect participants’ positive and negative moods. Moreover, an ANOVA on the post-pre differentials indicated no main effects of valence (*F*(1, 21) = 0.01, *p* = .927, *η*^2^_p_ < .001) and movement (*F*(1, 21) = 0.09, *p* = .766, *η*^2^_p_ = .004) and their interaction (*F*(1, 21) = 0.23, *p* = .634, *η*^2^_p_ = .011) on the PANAS differential score. These further suggest that mood changes were null or minimal, regardless of mood valence and the task conditions.

**Table 1.**
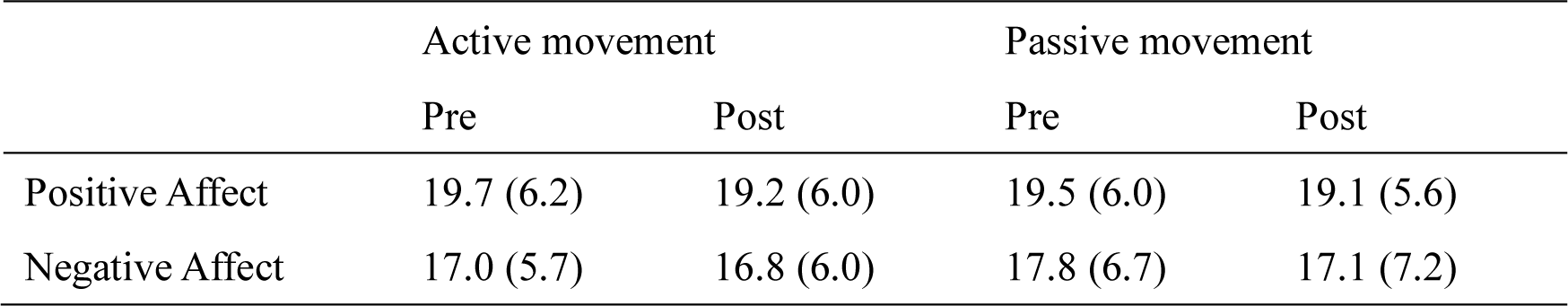
Mean scores on the subscales of the Positive and Negative Affect Schedule in Experiment 2 (*N* = 22). SD in parentheses.

As shown in Table 2, we did not find any significant correlation between the degree of intentional binding in each delay condition and changes in Positive Affect and Negative Affect under the active and passive conditions (|*r*|s < .332, *p*s > .131), again suggesting that intentional binding is not merely an artifact of mood changes induced by manual actions.

**Table 2.** Pearson’s correlation coefficients between the degree of intentional binding (i.e., difference between reproduction error in the passive and active conditions) and differentials of the Positive Affect (ΔPA) and Negative Affect (ΔNA) between post- and pre-task administrations in Experiment 2. *p*-values in parentheses (*df* = 20).

| | | | $\Delta PA$ | | $\Delta NA$ | |
| --- | --- | --- | --- | --- | --- | --- |
|  |  |  | Active | Passive | Active | Passive |
| Degree of intentional binding | 300-ms delay |  | -.212 (.343) | -.033 (.885) | -.323 (.143) | .183 (.416) |
|  | 400-ms delay |  | -.265 (.233) | .332 (.131) | -.095 (.674) | .022 (.921) |
|  | 500-ms delay |  | -.057 (.800) | .088 (.697) | .081 (.722) | -.040 (.861) |
|  | Averaged |  | -.223 (.318) | .149 (.508) | -.154 (.493) | .074 (.743) |

#### 3.2.3. Summary of Experiment 2

Results of Experiment 2 suggested that the time reproduction method is able to detect intentional binding in an action–outcome dyad. Active and passive manual movements did not induce substantial changes in emotional states (i.e., positive and negative moods), and if any, mood changes were unlikely to affect intentional binding. Thus, findings from Experiment 1 can be explained not merely by the artefacts of methodological and emotional influences, rather by the effect of active and passive movements *per se*.

## 4. General discussion

The present study investigated the intentional binding (i.e., a compression of subjective temporal interval between voluntary action and its sensory outcome) in more ecological settings, that is, alternations of the action–outcome dyad. Results of Experiment 1 suggested that the subjective duration of action–outcome alternations can be shortened when participants perform active finger movements than when they engage in passive movements. This finding extends the intentional binding effect, which has been robustly found in a single action–outcome dyad, and has been employed as an implicit measure of sense of agency. In the other session with identical setup, participants estimated the first and last action–outcome dyads within alternations. Contrary to the above intentional binding in the alternations, the perceived temporal interval of the first dyad was not affected by active or passive movement, although follow-up Experiment 2 suggested that the time reproduction method *per se* was able to detect intentional binding in situations in a single action–outcome dyad. Intriguingly, the perceived last dyad interval showed the opposite direction of temporal binding (i.e., *dilation* of perceived interval induced by active movement). Importantly, different biases in perceived durations and intervals could not be merely explained by the differences in the variability of time reproduction and mood changes between active and passive movements. Furthermore, intentional bindings for the action–outcome alternations and single dyad did not correlate, suggesting that the perceived duration of action–outcome alternations was not simply a summation of the perceived intervals of each action–outcome dyad. We propose that, in action–outcome alternations, different types of intentional binding (i.e., global compression and local dilation) might emerge depending on what individuals recall.

### 4.1. Compressed action–outcome alternations: An extended intentional binding

The compressed subjective duration of action–outcome alternations in conditions with active movements in contrast to those with passive movement can be considered as an extension of the intentional binding that has been observed robustly and extensively in an action–outcome dyad (Caspar et al., 2015; Engbert et al., 2008; Haggard et al., 2002). In line with previous studies on the intentional binding for an action–outcome–action triad (Yabe et al., 2017) and that for two outcomes (Ruess, Thomaschke, Haering, et al., 2018), our findings suggest that intentional binding could also occur in action–outcome alternations, which may simulate a more ecological situation, potentially implying social situations such as conversation and playing catch. Employing a mutual task performed by two participants and natural sensory outcomes (e.g., voice) in future studies could provide further insights into the social function of intentional binding.

One might argue that the subjective duration compression might not be a result of “active” movements but due to some artefacts. In our data, the variability (or stability) of time reproduction was comparable between active and passive movements, suggesting that the voluntariness and/or motor commands *per se* affected the perceived duration of action–outcome alternations. Furthermore, results from Experiment 2 suggested that participants’ emotional states, which affects the internal clock and consequently time perception (Droit-Volet, 2018; Droit-Volet et al., 2011) and intentional binding (Christensen et al., 2019), cannot merely explain the compressed subjective duration in the active-movement condition. Aside from the influence of repetition of active movements, our data also suggested a commonality between the empty and filled time during voluntary action. In most previous studies, intentional binding has been measured for an *empty* temporal interval between an action and its outcome. In contrast, participants in the present study reproduced a duration of stimulus (i.e., fixation cross) that was *filled* with actions and outcomes. It has been known that filled durations of sub- and supra-seconds are likely to be perceived to last longer than are empty intervals for the identical duration (Thomas & Brown, 1974). Moreover, this “filled-duration illusion” can be explained by a change in the internal clock speed (Wearden, Norton, Martin, & Montford-Bebb, 2007). Regardless of the known characteristics of filled duration, we observed intentional binding (i.e., compression) for the duration filled with actions and outcomes. Taken together, these findings could suggest that intentional binding may not simply stem from the basic time perception system that postulates the internal clock (for details see Section 4.3).

Increasing the temporal discrepancy between each action and its outcome violates the internal prediction of the sensory feedback corresponding to motor commands and can decrease the amount of intentional binding between a voluntary action and its outcome (Haggard et al., 2002; Imaizumi & Tanno, 2019; Ruess, Thomaschke, & Kiesel, 2018) and the intensity of the explicitly felt sense of agency over the outcome (Imaizumi & Tanno, 2019; Sato & Yasuda, 2005). Because of this similarity, intentional binding in an action–outcome dyad has been thought as an implicit measure of sense of agency. We showed that the amount of compression of subjective duration of action–outcome alternations is also decreased by outcome delays. Although the present experiment did not assess the explicit sense of agency over action and/or outcome, we speculate that the intentional binding in action–outcome alternations might also reflect the sense of agency during continuously performed action (Asai, 2017; Imaizumi & Asai, 2017).

### 4.2. Unaffected or even dilated action–outcome dyad in alternations

Perceived intervals of the first and last action–outcome dyads showed a tendency that was different from that observed in the perceived duration of action–outcome alternations exhibiting intentional binding. As for the first dyad, there was no difference between the active- and passive-movement conditions, suggesting no occurrence of intentional binding. Given that there is little evidence of intentional binding for intervals shorter than 500 ms as measured by the time reproduction method (but see Dewey & Knoblich, 2014), one might argue that the methodological artefact could nullify intentional binding. However, results of Experiment 2 suggested that the reproduction method for subsecond intervals *per se* is able to detect intentional binding. We speculate potential influences of serial position of the recalled interval in a sequence of intervals (e.g., action–outcome dyads). In our experiment, the first dyad interval was the most distant from the moment of recollection and reproduction. Memory of durations or intervals relies on mechanisms analogous to working memory (Manohar & Husain, 2016; Teki & Griffiths, 2014). For example, when individual is passively presented a sequence of different subsecond intervals and reproduces one of the intervals, the precision of the interval reproduction follows a tendency similar to the serial position effect (Murdock, 1962), where the first and last items are recalled better (i.e., primacy and recency effects, respectively). Therefore, the first and last intervals in a sequence are likely to be reproduced more veridically than are the other intervals (Teki & Griffiths, 2014). If we consider intentional binding (i.e., underestimation in the active movement condition) as a veridical response or typical performance, the expected intentional binding for the first dyad interval might have been nullified by a lack of primacy effect due to extinction and distraction by the subsequent intervals. However, we found that at least the time reproduction performance *per se* was not affected by serial position, as evident from the post-hoc ANOVA that showed no effect of serial position (first-versus last-dyad interval) on the reproduction error (*F*(1, 23) = 0.59, *p* = .449, *η*^2^_p_ = .025) or coefficients of variation (*F*(1, 23) = 1.41, *p* = .248, *η*^2^_p_ = .058).

On the other hand, temporal binding between the “outcome–action” dyad could be an alternative potential explanation. It has been known that the timing of a stimulus triggering an action can be perceived to be attracted toward the action, resulting in a compressed interval of the outcome– action dyad (Yabe et al., 2017; Yabe & Goodale, 2015). Given this, if the first tone served as a trigger for the subsequent (second) keypress, the subjective onset timing of the first tone might have shifted towards the subsequent keypress. This may result in the dilating effect on the first dyad interval and may compete against the known intentional binding for the first dyad, finally resulting in no apparent time distortion (i.e., no difference between active and passive conditions).

In contrast, the last dyad interval was perceived to last longer in the active than the passive condition, apparently suggesting an inverse direction of intentional binding. We again discuss this finding based on the potential role of stimulus triggering action (Yabe et al., 2017; Yabe & Goodale, 2015). For the last dyad, the preceding (third) tone might have attracted the subjective timing of onset of the subsequent (fourth) keypress, resulting in the dilated interval of the last keypress–tone dyad. However, we must be cautious when interpreting Yabe and colleagues’ findings, because they demonstrated shift in the perceived timing of the *trigger tone* towards the subsequent action. Furthermore, it should be explained why (expected) intentional binding for the last dyad was replaced by the potential effect of the trigger tone so clearly. It seems as if the perceived keypress timing chose the preceding tone as a direction of shifting. The cue integration theory posits that sense of agency is a result of combining internal and external cues (e.g., motor and sensory signals) weighted by their precisions in terms of information notifying the source of action and its outcome (Moore & Fletcher, 2012; Moore, Wegner, & Haggard, 2009). This theory can be applied to intentional binding (Moore et al., 2009; Wolpe, Haggard, Siebner, & Rowe, 2013), which can be considered as a marker of sense of agency. For example, an event which has relatively less reliable information is perceived to shift in time towards the other event with more reliable information. Specifically, the perceived timing of keypress can shift more strongly to the subsequent tone when the latter has a strong intensity against a background white noise (i.e., more reliable information) than when it has a weak intensity (Wolpe et al., 2013). In our task, the third tone preceding the fourth keypress might have served as an action trigger. Indeed, participants were instructed to make the keypress according to the preceding tone. Thus, we speculate that a tone triggering a keypress could have more reliable information as an external cue for temporal binding than would a non-trigger outcome tone, resulting in the temporal shift of the perceived timing of the keypress to the preceding trigger tone rather than to the outcome tone. Nevertheless, we should note another potential account for the last dyad dilation. Yabe et al. (2017) also found that there was no shift in perceptual timing of a tone between two keypresses. This suggests that the tone, which can serve as an action trigger and an action outcome, can shift in time towards the preceding and following keypresses although such bindings may cancel out each other. Hence, it can also be speculated that the fourth tone in our task had a prioritized role as an action trigger and thus its perceptual timing shifted towards the fifth keypress, resulting in the dilation of the last dyad interval.

### 4.3. Independent distortion of global and local time

Based on the scalar expectancy theory that postulates an internal clock counter producing the magnitude of perceived time (Gibbon, Church, & Meck, 1984; Wearden, 1991), one might expect that the intentional binding for the duration of the whole action–outcome alternations results from an accumulation of the local time distortion of each action–outcome dyad, such as (if any) the intentional binding for an action–outcome dyad. If so, the degree of intentional binding for the trial duration in Experiment 1 (i.e., difference between active and passive conditions) would correlate with that for the first and/or last dyad interval. However, we did not find any inter-participants correlation between these three measures, suggesting that the intentional binding for the action–outcome alternations and single action–outcome dyad could be independent even though they are experienced in virtually the same audiomotor task. This dissociation between the time perception of global and local events suggests a dedicated but not general internal clock that generates subjective time flow. Fereday and Buehner (2017) have performed a series of experiments to show that, while the perceived interval between an action and its outcome is robustly compressed, the perceived duration of a stimulus embedded in the interval remains unaffected. They concluded that a dedicated internal clock selectively induces the compression of the action–outcome interval. Our results also support the dedicated clock account, since the compression of duration of the action–outcome alternations did not necessarily entail the compression of the local action–outcome dyad interval.

In fact, the last dyad interval was perceived to dilate rather than compress in the active movement condition, potentially suggesting that different biases from the internal clock and cue integration could inherently prevail but independently affect the subjective time perceived during voluntary action based on what the individual intends to recall and reproduce. According to a recent “postdiction” framework (Shimojo, 2014), physical and sensorimotor events come into the brain and construct an implicit representation of the event sequence at early levels. At the later stage, a conscious percept is constructed using the previous sensorimotor information and even other sources so as to, for example, maintain the contextual consistency and retain the sense of agency. We speculate that, in our experimental task, the sensorimotor signals and temporal information generated through the action–outcome alternations might firstly generate an implicit representation of the duration/interval of the action–outcome alternations and single dyad. Then, when individuals recall and *postdict* an experienced time, the implicit representation of time may be reconstructed by a separate mechanism, such that the duration/interval they recall can be selectively distorted. For example, you may experience compression of the duration of action– outcome alternations only when you recall it.

### 4.4. Conclusions

The present study showed a subjective compression of the duration of alternating active movements and auditory outcomes as compared to a condition with passive movements (i.e., intentional binding in action–outcome alternations). Although we did not measure explicit sense of agency, the binding in alternating actions might provide insight into time perception and sense of agency in ecological and social contexts. In contrast to the “global compression,” the local action–outcome interval was unaffected or even dilated. The global and local levels of subjective time in the action–outcome alternations did not correlate, suggesting that our subjective time perception during action may rely not only on the general internal clock with scalar timing; rather, it would also be affected by other postdictive biases that are flexibly switched based on what we recall.

## Declaration of interest

The authors declare no conflict of interest.

## Funding

This work was supported by Grant-in-Aids for JSPS Research Fellow (16J00411) and Young Scientists (B) (17K12701) to SI, and Grant-in-Aids for Scientific Research on Innovative Areas ‘‘Understanding brain plasticity on body representations to promote their adaptive functions’’ (26120002) and for Scientific Research (B) (18H01098) to HI from the Japan Society for the Promotion of Science who had no involvement in the conduction of this study or manuscript preparation.

### Acknowledgements

The authors thank Yuta Kobayashi for his help during data collection in Experiment 1.

## Author contributions

SI conceived the study, performed the experiments, and analyzed the data. SI, YT, HI interpreted the results, wrote the manuscript, and approved the final version of the manuscript for submission.

